# YibN, a bona fide interactor of YidC with implications in membrane protein insertion and membrane lipid production

**DOI:** 10.1101/2024.12.13.628402

**Authors:** Zhiyu Zhao, Nachi Yamamoto, John W. Young, Nestor Solis, Amos Fong, Mohammed Al-Seragi, Sungyoung Kim, Hiroyuki Aoki, Sadhna Phanse, Hai-Tuong Le, Christopher M. Overall, Hanako Nishikawa, Mohan Babu, Ken-ichi Nishiyama, Franck Duong van Hoa

## Abstract

YidC, a prominent member of the Oxa1 superfamily, is essential for the biogenesis of the bacterial inner membrane, significantly influencing its protein composition and lipid organization. It interacts with the Sec translocon, aiding the proper folding of multi-pass membrane proteins. It also functions independently, serving as an insertase and lipid scramblase, augmenting the insertion of smaller membrane proteins while contributing to the organization of the bilayer. Despite the wealth of structural and biochemical data available, how YidC operates remains unclear. To investigate this, we employed proximity-dependent biotin labeling (BioID), leading to the identification of YibN as a crucial component within the YidC protein environment. We then demonstrated the association between YidC and YibN by affinity purification-mass spectrometry assays conducted on native membranes, with further confirmation using on-gel binding assays with purified proteins. Co-expression studies and *in vitro* assays indicated that YibN enhances the production and membrane insertion of YidC substrates, such as M13 and Pf3 phage coat proteins, ATP synthase subunit c, and various small membrane proteins like SecG. Additionally, the overproduction of YibN was found to stimulate membrane lipid production and promote inner membrane proliferation, perhaps by interfering with YidC lipid scramblase activity. Consequently, YibN emerges as a significant physical and functional interactor of YidC, influencing membrane protein insertion and lipid organization.

## Introduction

Membrane proteins (MPs) constitute approximately 20-25% of the cellular proteome and are essential for various functions, including signal transduction, nutrient transport, and energy production (1). Their proper assembly into the lipid bilayer is critical for maintaining membrane functionality. In *Escherichia coli*, the insertion of MPs into the membrane relies on the Sec translocon (2, 3). This machinery consists of the core SecYEG complex, which forms the membrane channel for protein translocation, along with additional factors such as the SecDFYajC sub-complex, the SecA ATPase, the YidC insertase and several other accessory components (4–7). Together, these components ensure the proper insertion and folding of MPs within the lipid bilayer, thereby upholding the integrity of the membrane bilayer in both form and function.

While YidC has a pivotal role in the biogenesis of multi-pass membrane proteins via the Sec translocon, it also functions in a Sec-independent manner to help the insertion of small membrane proteins (8, 9). YidC is a member of the conserved YidC/Alb3/Oxa1 insertase family, functioning within the bacterial, thylakoidal, and mitochondrial inner membranes, respectively, to facilitate the insertion of crucial components into energy-transducing complexes (10, 11). This includes the F1F0 ATP synthase subunit c (F0c) vital for ATP production. Additionally, in bacteria, YidC plays a role in the insertion of phage coat proteins M13 and Pf3, mechanosensitive channel subunits such as MscL, and components of the type VI secretion system like TssL (12–15). Despite the established importance of YidC in these processes, further research is required to elucidate the mechanisms by which the insertase facilitates the incorporation of small membrane proteins into the lipid bilayer.

Protein interactions and co-regulations offer critical insights into underlying molecular mechanisms, often explained by the guilt-by-association principle (16, 17). An illustrative example is the use of chemical cross-linkers, which revealed that the protease FtsH and its regulatory partners, HflC and HflK, are in proximity to YidC, thereby emphasizing the importance of protein quality control in YidC-mediated membrane insertion (18, 19). Investigation into YidC depletion also revealed a co-regulation of a specific glycolipid, termed MPIase (20). This glycolipid was subsequently shown to significantly enhance the insertion efficiency of various small membrane proteins, including F0c and phage proteins Pf3 and M13 (21, 22). Thus, knowledge of interaction networks can inform and help elucidate complex biological processes.

In this study, utilizing proximity-dependent biotin labeling (BioID) (23), we identify YibN, a 16 kDa single-pass inner membrane protein oriented towards the cytosol. We validate the YidC-YibN interaction *in vitro* and confirm its reliance on an intact N-terminal transmembrane segment. While YibN is not essential for cell viability, its expression significantly enhances the production of various small membrane proteins, including M13 procoat (PC-Lep), Pf3 (Pf3-23Lep), and F0c, as well as SecG. These latter findings are supported by *in vitro* protein translation and insertion assays. Additionally, the overproduction of YibN also correlates with increased membrane lipid synthesis and inner membrane proliferation and recent studies have indicated that lipid scrambling and bilayer reorganization are linked to membrane insertase activity (24, 25). These findings together place YibN as a bona fide interactor of YidC.

## Results

### Identifying potential interactors of YidC using BioID

The mutant biotin ligase BirA* (BirA R118G) (23) was fused to the C-terminus of YidC on plasmid pBAD22 (**Supplementary Figure 1**). After protein expression, the bacterial inner membrane was isolated and solubilized with 1% DDM. The extracted proteins were visualized on SDS-PAGE and those biotinylated were detected by Western Blot (**Figure 1A**). As expected, many proteins were biotinylated with YidC-BirA* compared to YidC-BirA, and none with YidC (right panel, **Figure 1A**). The detergent extract was also incubated with NeutrAvidin beads to isolate the biotinylated proteins. The eluted proteins were trypsin-digested and analyzed by LC-MS/MS. A total of 172 proteins were detected across four replicates, with 21 proteins found in common across the replicates (**Supplementary File 1**). The proteins identified were ranked based on their spectral counts and plotted in **Figure 1B**. As expected, YidC and BirA were efficiently detected in the samples. The protein BCCP (gene name: *accB*), the natural substrate of BirA, was also identified with a high spectral count. Other proteins, such as FtsH, HflK, and HflC, previously reported vicinal to YidC based on chemical crosslinking analysis (18), were also efficiently detected. Strikingly, the protein with the highest spectral counts, and consistently across four replicates, was the uncharacterized protein YibN.

**Figure 1.**
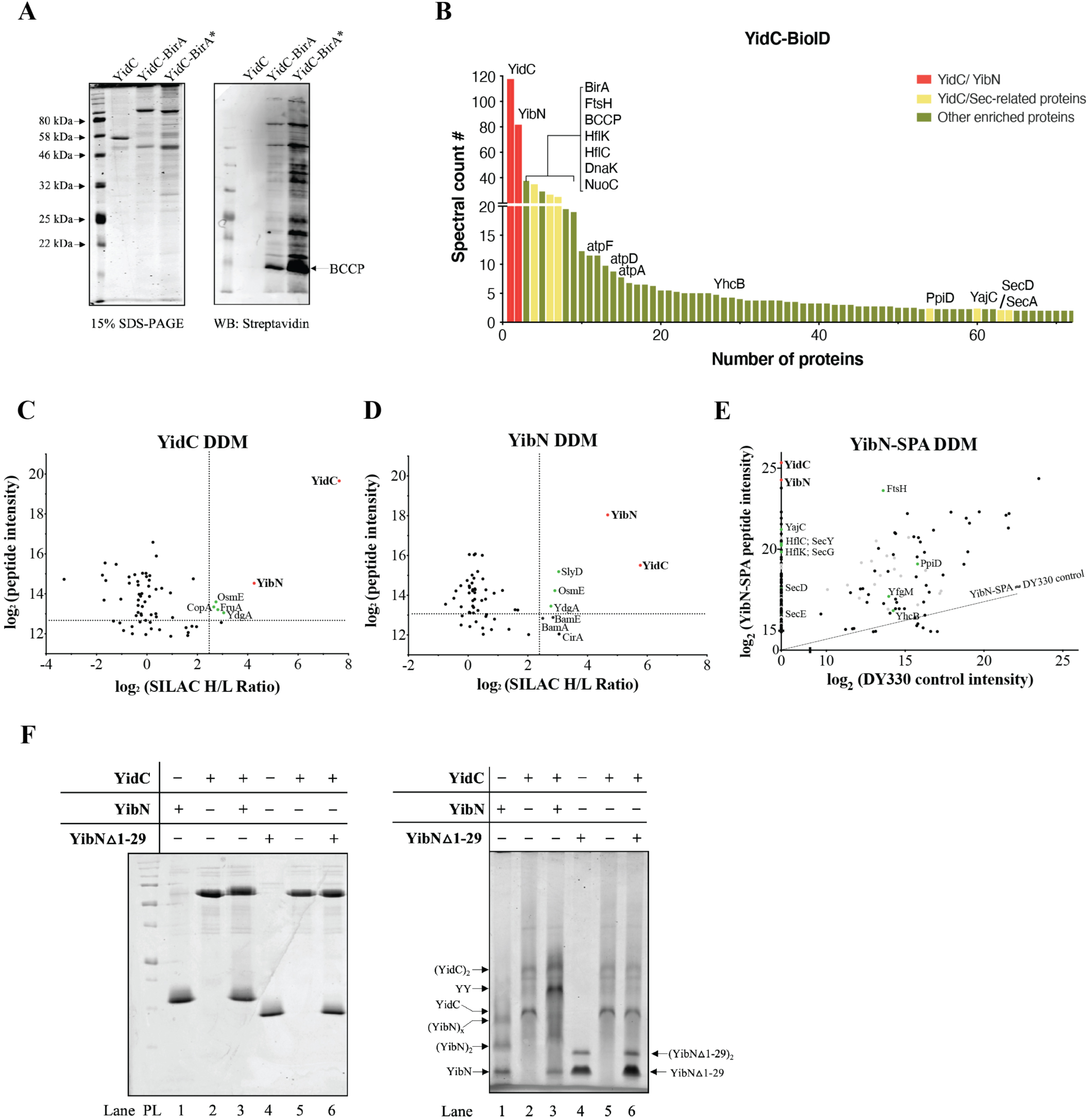
Identification and validation of the YidC-YibN interaction. **(A**) BirA* was fused to the C-terminus of YidC. Expression of the fusion protein YidC- BirA* allows biotinylation in *E. coli*. Following the inner membrane vesicle (IMV) isolation and solubilization in detergent DDM, the biotinylated proteins were pulled down via streptavidin resin. The eluted proteins are analyzed on SDS-PAGE and Western Blot. The aliquots of IMVs of YidC-BirA* and its controls YidC-only and YidC-BirA are loaded onto 15% SDS-PAGE followed by Coomassie staining. The biotinylation was verified on Western Blot (Streptavidin anti-biotin). **(B**) Top hits of YidC-BioID identified by LC-MS/MS. YidC and YibN are highlighted in red. YidC/Sec-related proteins (reported interacting with YidC or in the vicinity of YidC: van Bloois *et al.*, 2008; Sachelaru *et al.*, 2017; Young *et al.*, 2022) are labeled in yellow. “Other enriched proteins” are labeled in green. **(C-D)** Validation of YibN-YidC interaction by AP-MS. SILAC-AP/MS was conducted in detergent DDM. The crude membranes containing either (**C**) YibN-His6 or (**D**) His6-YidC obtained from Lys4-labeled *E.coli* JW2806 strain were solubilized in 0.8% DDM followed by Ni-NTA purification. Empty vector pBAD33 or pBAD22 were used as controls and expressed in Lys0-labeled media. The log2(peptide intensity) of each protein identified in the pull-down assay was plotted against its corresponding log2(SILAC H/L ratio). The proteins with an H/L ratio larger than 5 (log2(SILAC H/L ratio) > 2.4) will be considered “enriched”. The y-axis cutoff was set to the log2(peptide intensity) point, where three-quarters of data points will be above this cutoff. YidC/YibN are highlighted in red. Proteins with log2(SILAC H/L ratio) larger than 2.4 and top 75% abundant proteins are labeled in green. **(E)** Endogenous YibN-SPA pull-down assay. *E. coli* strain B3611 bearing SPA-tagged YibN was grown. The pull-down assay was conducted in detergent DDM. YidC and YibN are labeled in red. Known interactors of YidC (van Bloois *et al.*, 2008; Sachelaru *et al.*, 2017; Young *et al.*, 2022) are labeled in green. Ribosomal proteins are labeled in grey. Proteins between YibN and YajC are GntR, YgfX, QmcA, and CorA. **(F)** Biochemical evidence of YidC and YibN complex. YidC, YibN, and YibN△1-29 sample analysis on SDS-PAGE (containing β-ME), followed by Coomassie Blue staining (left panel). YidC-YibN binding analysis on BN-PAGE. 3 µg of YibN, YidC or YibN △1-29 were loaded as indicated.

### Validation of YibN-YidC interaction using affinity pulldown

Recombinant His-tagged YidC and control empty plasmid pBAD22 were transformed into strain JW2806 to allow SILAC-labeling with Lys4/Lys0 lysine isotopologues. After YidC expression, the membrane fraction was solubilized with DDM and incubated with Ni-NTA agarose beads. After washes, the proteins retained with YidC were identified by LC-MS/MS (**Figure 1C**). As expected, His-tagged YidC was well detected in the sample, with >150-fold enrichment over the background. Strikingly, endogenously expressed YibN was also detected, with >20-fold enrichment over the background. In the reciprocal experiment, His-tagged YibN and the control empty plasmid were transformed into strain JW2806.

Recombinant YibN was enriched in the sample (>20-fold) and endogenous YidC was also well detected (>50-fold over the background; **Figure 1D**). The pulldown experiments were then conducted with the recombinant strain B3611 (26). In this strain, the chromosomally encoded YibN protein is tagged at the C-terminus with a peptide affinity SPA tag (16, 27). The untagged parental strain DY330 was employed as a control to subtract non-specific binding to the FLAG resin. The data, plotted in **Figure 1E**, show that YidC and YibN are the two most abundant proteins isolated from strain B3611 under native expression conditions.

### Validation of the YidC-YibN interaction by native-gel electrophoresis

His-tagged YidC and His-tagged YibN were purified to homogeneity in DDM detergent. The proteins were analyzed by SDS-PAGE (left panel, **Figure 1F)** and blue-native PAGE (right panel, **Figure 1F**). The blue-native gel analysis revealed that YidC and YibN form a distinctive band when incubated together (labeled YY in **Figure 1F**). To confirm the identity of this band, the YibN transmembrane segment (TMS; residue 1-29) was deleted. On the blue-native PAGE, the truncated △1-29 YibN failed to associate with YidC, indicating that the TMS of YibN is essential for the formation of the YidC-YibN complex.

### YibN overexpression leads to inner membrane proliferation

During the YibN purification procedure on a sucrose gradient, we noticed that a larger amount of inner membrane is produced compared to the control plasmid strain (left panel, **Figure 2A**). Total membrane lipids were extracted and analyzed by thin-layer chromatography (TLC; right panel, **Figure 2A**). This analysis confirmed that the YibN overproducing strain produced ∼4-fold more membrane lipids than the control strain, with phosphoethanolamine (PE) and phosphoglycerol (PG) remaining the predominant species.

**Figure 2.**
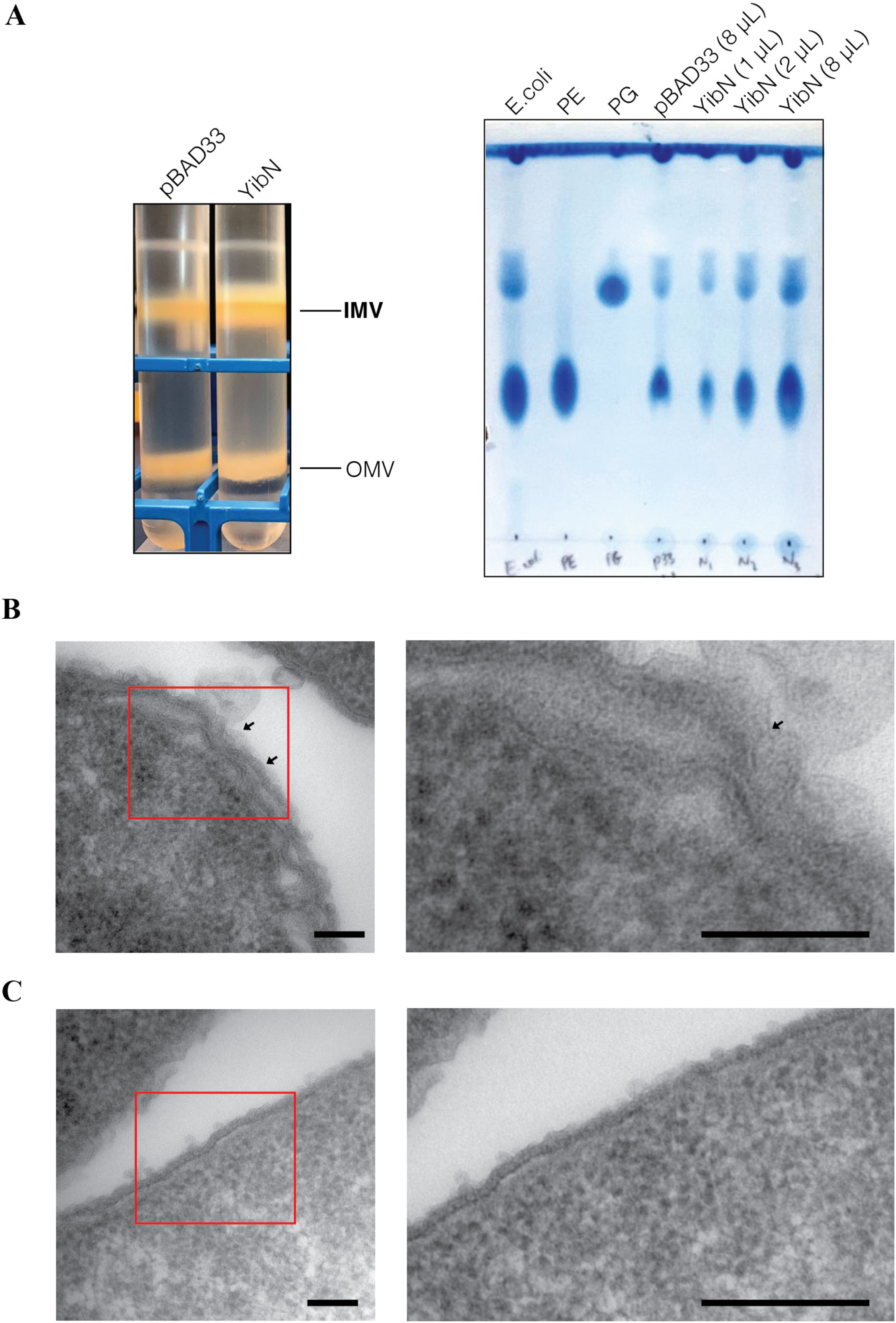
Membrane lipid analysis following YibN over-expression. **(A**) Inner and outer membrane isolation on sucrose gradient and lipid content analysis on thin layer chromatography (TLC). Left panel: IMV and OMV isolation of cells bearing either empty pBAD33 or pBAD33 YibN on 20-50-70% sucrose gradient. Right panel: TLC analysis of lipid extracts obtained from 30 µg of YibN overproduced inner membrane and 30 µg of control inner membrane. **(B**) Transmission electron micrographs of sectioned *E.coli* BL21 (DE3) with over-expressed YibN. BL21 (DE3) bearing YibN was grown at room temperature until OD600 reached 0.4. 0.1% arabinose was used to induce the overproduction of YibN. Cells continued to grow for another 1.5 hours. **(C)** A negative control, in which an empty vector was transformed and cells were grown and induced under the same condition as shown in (B). *Scale bar*, 100 nm. The right panels are the enlarged views of the area circled in red from the left panels.

The same cells were also stained and visualized by transmission electron microscopy. The images show that YibN production is associated with membrane proliferation, circumvolutions and multilayered structures principally at the level of the bacteria’s inner membrane (compare **Figure 2B** to **Figure 2C**). The overall shape of the outer membrane seems less affected (**Supplementary Figure 2**).

### YibN augments the biogenesis of YidC substrates

We tested the effect of YibN on three known YidC substrates: M13 procoat (PC) and Pf3 coat proteins, each fused to the soluble domain of leader peptidase (Lep) to facilitate their detection (28, 29), and F1-F0 subunit F0c (13). These substrates were co-expressed with YibN or the control empty plasmid pBAD33 (**Figure 3**) at room temperature using 0.1% arabinose for 15 minutes, followed by the addition of 0.75 mM IPTG to induce substrate synthesis.

**Figure 3.**
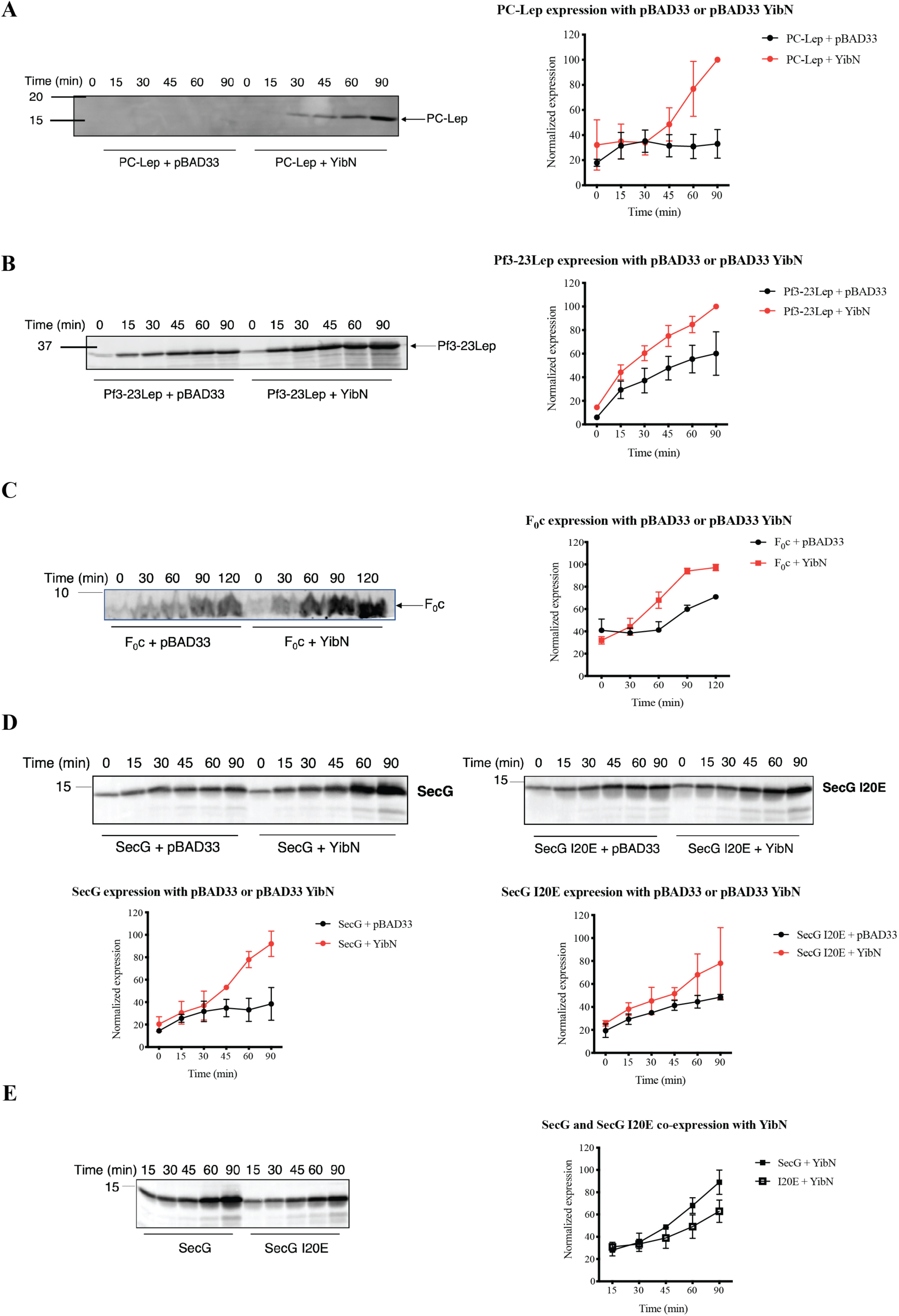
Effect of YibN on small membrane proteins biogenesis. (A-D) PC-Lep, Pf3-23Lep, F0c or SecG co-expression with empty pBAD33 or pBAD33 YibN respectively at room temperature. The whole-cell lysates were analyzed by Western Blot (PC-Lep and Pf3-23Lep blots: anti-Lep antibody; F0c blot: anti-F0c antibody; SecG blot: anti-SecG antibody). The expression level of these small membrane proteins was quantified using ImageJ. The data represents the mean ± S.D. from the three independent experiments. **(E)** SecG WT and mutant I20E expression level comparison upon YibN overexpression. The SecG or mutant SecG level is quantified by ImageJ.

Aliquots were collected every 15 minutes for analysis by SDS-PAGE and Western Blot with the corresponding detection antibodies. The overall bacterial growth rate and cell protein content remained unchanged when the substrates were co-expressed with YibN or the empty plasmid (**Supplementary Figure 3A, 3B, 3C**). However, the synthesis of PC-Lep, Pf3-23Lep, and F0c was significantly increased in the presence of YibN (**Figure 3A, 3B, 3C**).

These co-expression experiments were also conducted with three other small membrane proteins, SecG, YajC, and YhcB (**Figure 3D, Supplementary Figure 3D, Supplementary Figure 4**). The biogenesis of the single-pass YajC and YhcB is not affected by YidC depletion (30), and it is unknown if SecG, which is topologically similar to F0c with two TMS, depends on YidC. Similarly, the control and experimental groups grew at similar rates and their overall cellular contents were comparable (**Supplementary Figure 3D**). The biogenesis of YajC and YhcB was not affected by YibN (**Supplementary Figure 4**), while the biogenesis of SecG was significantly increased (left panel, **Figure 3D**). Interestingly, the production of SecG carrying the mutation I20E in the first transmembrane segment was less affected by YibN (compare left and right panels in **Figure 3D**, and side-by-side comparison in **Figure 3E**), indicating that the hydrophobicity of TMS can be important in detecting the YibN effect.

### YibN stimulates protein insertion *in vitro*

To validate the YibN effect, we turned to an *in vitro* translation/insertion assay using inverted membrane vesicles (INVs) (31). Compared to the INVs prepared from the control strain, INVs enriched for YibN supported a 1.5∼1.8-fold stimulation of insertion of the substrates Pf3 coat, M13 procoat H5 and F0c (**Figure 4**). No such stimulation was obtained with INVs enriched for YidC, as previously reported (22). We also employed the SecG and SecG I20E protein substrates. The SecG protein is made with two TMS with N- and C-termini exposed to the periplasm, but SecG also exists in an inverted orientation, with its N- and C-termini exposed to the cytoplasm (**Supplementary Figure 5**) (32). After membrane insertion and proteinase K digestion, three membrane-protected fragments (MPF) were detected (*i.e*. MPF 1 and MPF 2 derived from normal SecG and one corresponding to the inverted SecG; **Figure 4D-E**). Their quantitation showed MPF 1, MPF 2, as well as inverted SecG, are all augmented with INVs enriched for YibN, as much as that obtained using INVs enriched for YidC (**Figure 4D**). In contrast, with the SecG I20E substrate, the stimulation was much less evident (**Figure 4E**).

**Figure 4.**
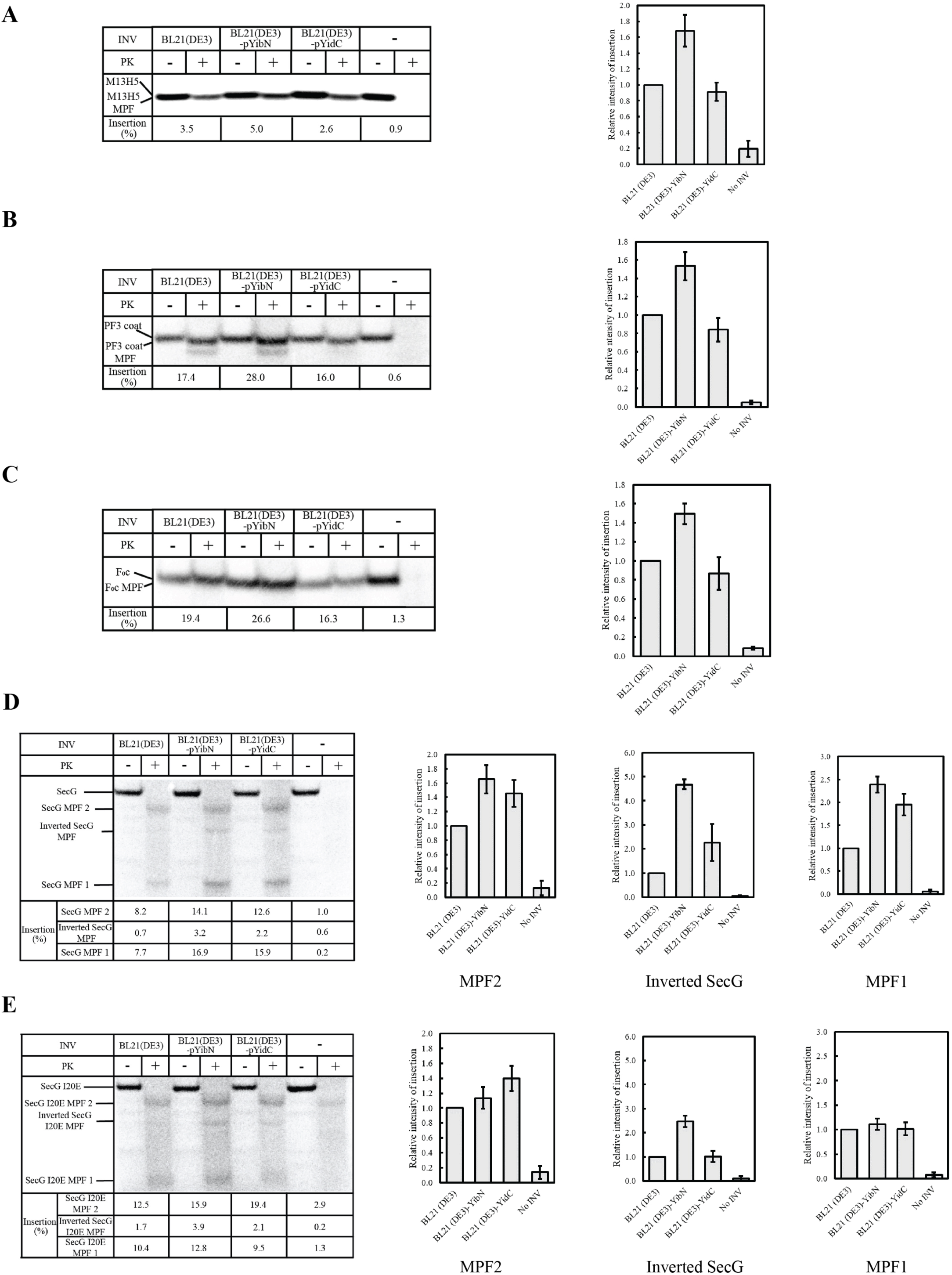
*In vitro* insertion assays of the small membrane proteins. M13 procoat H5 (**A**); Pf3 coat (**B**); F0c (**C**); SecG (**D**) and I20E SecG (**E**) were *in vitro* synthesized in the presence of wild-type INV (BL21(DE3)), YibN-overproduced INV (BL21(DE3)-pYibN) or YidC-overproduced INV (BL21(DE3)-pYidC) as described in “Materials and methods”. The control sample without INV was also set (‘-’ and ‘No INV’). The membrane-inserted domain was generated by proteinase K (PK) digestion as a membrane-protected fragment (MPF). The insertion efficiency was calculated and expressed as a percentage of the translated level. The experiments were repeated at least three times. The relative activity of insertion to that in wild-type INV was shown with the SD values in graphs. Note that the full-length F0c is given upon membrane insertion with PK digestion. In the case of SecG and I20E SecG, three kinds of MPF (MPF2, inverted SecG, MPF1) are generated with PK digestion. Their activities relative to those in wild-type INV are shown with the SD values. Positions of each protein and MPFs are indicated.

## Discussion

YidC belongs to the “Oxa1 superfamily”, which includes the closely related Oxa1/Alb3 proteins and the functionally analogous EMC3, TMCO1, GET1, and Oxa1L insertion factors (33–36). These proteins typically feature a conserved structure characterized by a membrane-exposed hydrophilic groove that facilitates the translocation of membrane proteins into the lipid bilayer (37, 38). This structural groove is linked to a membrane bilayer thinning mechanism that probably reduces the energy expenditure required for the translocation process (39, 40). Additionally, this groove is implicated in inter-leaflet membrane lipid scramblase activity, a trait that also characterizes the membrane insertase family (25). This dual functionality highlights the importance of these proteins have in maintaining membrane integrity while assisting in the proper insertion and arrangement of other membrane proteins (34, 41, 42).

In *E. coli*, the common view is that YidC possesses the structural elements enabling the protein to operate alone, although recent reports on the closely related Oxa1L identify TMEM126A as a specific interactor (36). In this study, we used BioID to probe the protein environment of YidC and found YibN - a previously uncharacterized inner membrane protein - to be highly proximal. Our SILAC-AP/MS experiments confirmed that YidC and YibN reciprocally capture each other (**Figure 1C, 1D**), with endogenous YibN enabling the enrichment of endogenous YidC (**Figure 1E**). To validate this further, we purified these two proteins and applied an on-gel binding assay to find that YidC and YibN form a stable complex in detergent (**Figure 1F**). The deletion of the YibN unique transmembrane segment abolished this association, implying that the hydrophobic region is critical for the stability of this complex and suggesting that the binary interaction likely occurs within the hydrophobic interior of the lipid bilayer. Collectively, the evidence supports the conclusion that YibN is a bona fide interactor of YidC, pointing to a possible role in membrane protein biogenesis.

YibN is a 16 kDa protein composed of an N-terminal transmembrane segment and a cytosolic rhodanese-like domain. Interestingly, it has been observed that YibN is upregulated upon the depletion of YidC or the depletion of SecDF-YajC (30, 43), suggesting a possible involvement of YibN in the membrane protein biogenesis pathway. Additionally, the gene *yibN* is located on the same operon as *grxC*, *secB* and *gpsA* (44), which encode for proteins with physiological relationships. The protein GrxC promotes protein disulphide bond reduction, SecB maintains proteins competent for secretion, and GpsA is involved in glycerophospholipid synthesis (45–50). YibN, however, is not essential in *E. coli;* the knockout strain JW3586 does not display any obvious growth defects at the different temperatures tested (**Supplementary Figure 6**). This non-essentiality complicates the analysis since without evident defects, distinguishing the physiological and biochemical contributions of YibN becomes more challenging.

In our co-expression experiments, we observed that the production of YibN increases the production of the YidC substrates PC-Lep, Pf3-Lep, and F0c (**Figure 3**). YibN also augmented the production of SecG, but not when SecG carrying the mutation I20E located in its first transmembrane segment (**Figure 3D**). Two other small membrane proteins, YajC and YhcB, whose production is not affected by the YidC depletion (19) were not affected by YibN (**Supplementary Figure 4**). As additional experimental support, we turned to an *in vitro* translation and insertion assay (**Figure 4**). The substrates PC-Lep, Pf3-23-Lep, F0c and SecG, but not SecG I20E, all showed significant increased insertion using membranes enriched with YibN, whereas membranes enriched with YidC had less effect (**Figure 4**).

Studies have shown that YidC-dependent translocation is contingent on the MPIase glycolipid such that YidC overexpression would have less bearing on translocation insofar as MPIase level content remains constant (21, 22).

Interestingly, YibN expression induced noticeable membrane lipid production alteration and local cell surface deformation (**Figure 2, Supplementary Figure 2**). Studies have found that cells lacking membrane insertases exhibit defects in membrane morphology and lipid metabolism, which align with their role as glycerophospholipids scramblases (24, 25, 51, and references within). We speculate herein that the overproduction of YibN could interfere with the YidC lipid transport activity. This interference may occur due to an imbalance in protein-protein interactions, potentially leading to the alteration of YidC’s ability to scramble lipids. The mechanistic basis that underlies YibN’s role in modulating YidC activity remains to be explored.

## Materials and Methods

### Strains and plasmids

The strains and plasmids used in this study are listed in **Supplementary Table 1**. Strain BL21 (DE3) was used for protein expression. All the constructs were made through the PIPE cloning method (52). The pAD22-HisYidC expression vector was obtained from our laboratory collection (53). The BirA gene was cloned from the *E. coli* genome into the pET28a vector and a point mutation (R118G) was introduced to obtain BirA*. The pBAD22 YidC-BioID construct was obtained by fusing BirA* onto the C-terminus of His6-YidC. All sequences (BirA, BirA*, and YidC-BioID) were confirmed through Sanger Sequencing.

### Growth of YidC-BioID and membrane preparation

A single colony-bearing pBAD22 YidC-BirA* plasmid was inoculated in a 10 mL overnight culture in LB media supplemented with 100 µg/mL ampicillin. The overnight culture was back-diluted 1/100 into 1 L fresh LB media supplemented with ampicillin and grown until OD600 ∼ 0.4. Protein expression was induced with 0.2% arabinose for another 2 hours. Cells were harvested by centrifugation (6,000 x g, 10 minutes) and resuspended in the TSG buffer (50 mM Tris pH 7.9, 100 mM NaCl, 10% glycerol) containing 1 mM PMSF. The cell lysate was obtained by passing the cells over a Microfluidizer (12,000 psi, three times). Unlysed cells were removed by centrifugation (3,000 x g, 10 minutes). The crude membrane fraction was obtained by ultracentrifugation (100,000 x g, 45 minutes) in a Ti45 rotor. The crude membrane pellet was resuspended in 10 mL of TSG buffer. The inner and outer membrane fractions were further isolated through a step-sucrose gradient (20-50-70%) as previously described (54). The inner membrane fraction was collected from the 20-50% sucrose interface. The inner membrane fraction was diluted with TSG buffer and centrifuged again to remove the sucrose (TLA-110 rotor, 164,000 x g, 20 minutes). The inner membrane fraction was resuspended in the TSG buffer.

### Biotinylated proteins enrichment assay

The YidC-BirA* inner membrane fraction (2 mg) was solubilized with 1% n-dodecyl-β-d- maltoside (DDM) for 1 hour at 4℃ with gentle shaking. The insoluble material was removed by centrifugation (100,000 x g, 15 minutes). The DDM-extracted material was incubated with 120 µL NeutrAvidin resin equilibrated in the TSG buffer containing 0.02% DDM. The sample was then split into four fractions (each fraction containing ∼500 µg of solubilized material plus 30 µL NeutrAvidin resin). Fractions 1 and 2 (technical replicates) were washed with 1x PBS buffer (137 mM NaCl, 2.7 mM KCl, 10 mM Na2HPO4, 1.8 mM KH2PO4) (eight times, 500 µL each) with 0.02% DDM. Fraction 3 was washed with 1 M NaCl (four times, 500 µL each) supplemented with 0.02% DDM and then 1x PBS + 0.02% DDM buffer (four times, each time 500 µL). Fraction 4 was washed with 10% DMSO (four times, each time 500 µL) and then 1x PBS buffer (four times, each time 500 µL). The biotinylated proteins were eluted with a solution containing 80% acetonitrile, 0.2% trifluoroacetic acid (TFA), and 0.1% formic acid. The samples were then dried in SpeedVac and resuspended in 100 µL of 20 mM NH4HCO3 before trypsin digestion. The tryptic peptides were then Stage-tipped and analyzed on LC-MS/MS.

### Expression of YidC and YibN in SILAC labeling conditions

Plasmids pBAD22, pBAD33, pBAD22 His6-YidC and pBAD33-YibN-His6 were transformed into *E. coli* JW2806 strain. SILAC labeling was performed as previously described (17).

Briefly, cells were grown overnight in M9 media supplemented with either 0.3 mg/mL Lys4 (^2^H4-lysine, “heavy” pBAD22 His6-YidC and pBAD33 YibN-His6; the experimental group) or 0.3 mg/mL regular lysine Lys0 (regular lysine, “light” for pBAD22 and pBAD33; the control group). On the second day, cultures were back-diluted 1/100 into fresh M9 media supplemented with either Lys4 (experimental group) or Lys0 (control group). The cells were grown at 37 ℃ until OD_600_ ∼ 0.4 and induced with 0.02% arabinose. After growth at room temperature overnight (∼16 hours), cells were harvested by centrifugation (3,000 x g, 10 minutes). The cell pellets were resuspended in 4 mL of TSG buffer containing 1 mM PMSF. Following cell lysis on a French Press (500 psi, three passes), unlysed cells were removed through centrifugation (3,000 x g, 10 minutes), and the supernatant was subject to ultracentrifugation (100,000 x g, 20 minutes) to obtain the membrane fraction.

### Isolation of SILAC-labeled YidC and YibN

Membranes prepared from the “heavy” (pBAD22 His6-YidC and pBAD33 YibN-His6) and “light” (pBAD22 and pBAD33) conditions were solubilized with 0.5% DDM in TSG buffer at 4℃ with gentle shaking. The insoluble material was removed by ultracentrifugation (100,000 x g, 15 minutes) and the detergent-solubilized material (∼500 µg each) was incubated with Ni-NTA agarose beads at 4℃ for ∼2 hours with gentle shaking. The resin was washed with the TS/DDM buffer (50 mM Tris, 100 mM NaCl, 0.02% DDM; 5 times, each time 1 mL). The proteins were eluted in TS/DDM buffer supplemented with 600 mM imidazole. The eluted proteins from the “heavy” pBAD22 His6-YidC and “light” pBAD22 samples were pooled together. The eluted proteins from the “heavy” pBAD33 YibN-His6, and “light” pBAD33 were pooled together. To remove detergents before MS analysis, the pooled samples were treated with 1 mL 100% ice-cold acetone and incubated overnight at-20℃.

The proteins were sedimented by centrifugation (16,100 x g, 10 minutes). The acetone was removed and the protein pellets were air-dried. The pellets were resuspended in 50 µL of 20 mM NH4HCO3 before trypsin digestion.

### YibN-SPA membrane isolation and FLAG pull-down

The procedures for YibN-SPA cell growth, membrane isolation and membrane solubilization were as described in Young *et al.*, 2022 (7). Briefly, strain B3611 (chromosomal expression of SPA-tagged YibN) and strain DY330 (control) were revived on LB-agar plates with/without Kanamycin antibiotic. A single colony was inoculated for overnight growth and back-diluted 1/100 into fresh LB media at 37℃ until the OD600 ∼ 1. Cells were harvested and lysed as described above. The membrane fraction was isolated by ultracentrifugation (100,000 x g, 30 minutes) followed by resuspension in the TSG buffer. The membrane fraction was solubilized in 0.5% DDM. The detergent extract (∼1 mg) was incubated with 50 µL of ANTI-FLAG M2 affinity gel at 4 °C overnight with gentle shaking. The resin was washed with TSG-DDM buffer (50 mM Tris, pH 7.8; 100 mM NaCl; 10% glycerol; 0.02% DDM) 6 times. Bound proteins were eluted in 100 mM Glycine HCl, pH 3.5. The pH was then adjusted to 8.0 with 1 M Tris, pH 8. The proteins were precipitated in acetone to remove detergent, as described above.

### MS sample preparation and analysis

The samples described above (YidC-BioID, YidC-SILAC-AP, YibN-SILAC-AP, and YibN- SPA) were treated with 6 M urea for 30 minutes at room temperature. Samples were treated with 10 mM DTT for 1 hour and with 20 mM IAA for another 30 minutes. 10 mM DTT was further added to the samples for 30 minutes. The samples were then diluted 5 times with the TS buffer (50 mM Tris, pH 7.8; 100 mM NaCl) and trypsin-Gold was added at an enzyme-to-protein ratio of 1:50 for 16 hours. The tryptic peptides were Stage-tipped and dried in SpeedVac. Data were acquired on a Bruker Daltonics Impact II quadrupole time-of-flight mass spectrometer connected online to an easy nLC-1000 HPLC (Thermo Scientific). A one-column set up was utilized and peptides were separated using a 35 cm, 75 µm I.D. column packed with 1.8 µm C18 resin (Dr Maisch). Peptides were loaded onto the column using buffer A alone (0.1% formic acid in water) at 800 bar for a total of 5 µl volume washed and then eluted on a linear gradient from 2% buffer B (0.1% formic acid in acetonitrile) to 35% buffer B in 60 min then washed with 90% buffer B for 10 min. The Impact II mass spectrometer was operated in data-dependent acquisition mode with a top20 method. MS1 scans were measured at 5Hz and collision-induced dissociation (CID) performed on the 20 most intense ions between charge states 2 and 6. Each MS/MS event was done at a scan rate of 20 Hz and excluded for 30 seconds unless the intensity of the precursor was 4x greater than its previous MS/MS event. Data were then processed with DataAnalysis to convert files into.mgf files.

All MS/MS samples (mgf files) were analyzed using Mascot (Matrix Science, London, UK; version 2.5.1). Mascot was set up to search the *Escherichia coli* K12 (accessed May 12 2016) database (8612 entries) assuming the digestion enzyme trypsin. Mascot was searched with a fragment ion mass tolerance of 0.100 Da and a parent ion tolerance of 50 PPM. Carbamidomethylation of cysteine (+57.021464 Da) was specified in Mascot as a fixed modification. Oxidation of methionine (+15.99491 Da) and biotinylation of lysine and the N-terminus (+226.077598 Da) were specified in Mascot as variable modifications.

Scaffold (version Scaffold 4.8.5, Proteome Software Inc., Portland, OR) was used to validate MS/MS based peptide and protein identifications. Peptide identifications were accepted if they could be established at greater than 98.0% probability to achieve an FDR less than 1.0%. Peptide Probabilities from Mascot were assigned by the Scaffold Local FDR algorithm.

Protein identifications were accepted if they could be established at greater than 84.0% probability to achieve an FDR less than 1.0% and contained at least 1 identified peptide. Protein probabilities were assigned by the Protein Prophet algorithm (55). Proteins that contained similar peptides and could not be differentiated based on MS/MS analysis alone were grouped to satisfy the principles of parsimony. Spectral counts for identified proteins were determined through Scaffold and used to estimate relative abundance of proteins.

Quantitative analysis of SILAC labelled samples: Raw files from the Impact II MS system (.d folders) were searched directly with Maxquant. Quantitative modifications used ^15^N labeling for analysis, fixed modification of carbamidomethylation of cysteine (+57.021464 Da), and variable modifications: oxidation of methionine (+15.99491 Da) and biotinylation of lysine and the N-terminus (+226.077598 Da) (if for BioID experiment). Peptide and protein FDRs were set to 1% and results later imported and analyzed in the Perseus software environment.

### YidC, YibN and YibN△1-29 purification and binding assay

Plasmids pBAD22 His6-YidC, pBAD33 YibN-His6, and pBAD33 YibN△1-29-His6 were transformed into BL21 (DE3) strain. The procedures to grow cells and isolate membranes of His6-YidC and YibN (full length) were as described above (section “Growth of YidC-BioID and membrane preparation”). The protein expression was verified by loading aliquots of the membrane and soluble fractions on a 15% SDS-PAGE followed by Coomassie blue staining. Membranes were solubilized with 1% DDM for 1 hour at 4℃ with gentle shaking. The insoluble material was removed by centrifugation (100,000 x g, 15 minutes). The detergent extract or cytosolic fraction (∼ 8 mg each) was passed through a Ni-NTA gravity column (1 mL resin). The flowthrough was collected, and the resin was washed with 50 mL TSG-DDM buffer plus 20 mM imidazole. Bound proteins were then eluted with TSG-DDM buffer supplemented with 600 mM imidazole. Purified YidC (3 µg) was incubated with either purified YibN or purified YibN △1-29 (3 µg each) for 15 minutes on ice. The protein samples were then analyzed by Blue Native-PAGE and 15% SDS-PAGE.

### BW25113 and JW3586 growth at various temperatures

A single colony of strains BW25113 (no antibiotic) and JW3586 (25 µg/mL Kanamycin) was inoculated and grown in LB media overnight, respectively. On the second day, based on the OD600 reading, a similar number of cells were back diluted 1/50 into fresh LB media with no antibiotics. The cells were grown at 18℃, 37℃ or 42℃ for 3 hours. Later, two strains were plated on LB agar plates and the cells were recovered at 37℃. A serial dilution with a dilution factor of 10 was performed.

### Co-expression of YibN with the YidC substrates PC-Lep, Pf3-Lep, F0c and SecG

Plasmids pMS119 PC-Lep, pMS119 Pf3-Lep, pET20 *atpE* (encoding F0c) and pBAD22 SecG were co-transformed with either empty vector pBAD33 (control) or pBAD33 YibN plasmid (experimental group) into strain BL21 (DE3). For each co-transformant, a single colony was inoculated into 10 mL LB media plus 10 µg/mL Ampicillin and 30 µg/mL Chloramphenicol antibiotics and grown overnight at 37℃. On the second day, the overnight cultures were back diluted 1/100 into fresh LB media with antibiotics. The cells were grown at room temperature until their OD600 reached 0.4. For the co-expression of pBAD22 SecG with empty pBAD33 or with pBAD33 YibN, 0.1% arabinose was used for protein induction. The cells continued to grow for another 1.5 hours. For the co-expression of pMS119 PC-Lep, pMS119 Pf3-Lep, and pET20 *atpE* with empty pBAD33 or with pBAD33 YibN, the cells were firstly induced with 0.1% arabinose for 15 minutes. The cells were then induced with 0.75 mM Isopropyl β-D-1-thiogalactopyranoside (IPTG) for another 1.5-2 hours. Time-course aliquots were taken and analyzed by SDS-PAGE and Western Blot.

### *In vitro* membrane protein insertion assay

BL21(DE3) and BL21(DE3) carrying plasmid pBAD33 YibN or pBAD22 YidC were cultivated in LB medium at 25°C. At the mid-log phase, 0.2% arabinose was added to express YibN and YidC, respectively. After cultivation was continued for 3 h, INV (inverted membrane vesicles) were prepared as described (56) and then subjected to the insertion assay *in vitro*. The reaction mixture (20 μL) was composed of the pure system, a reconstituted translation system (57), INV (0.5 mg protein/mL), plasmid DNA in which the substrate membrane proteins are encoded under the control of the T7 promoter (50 μg/mL), and [^35^S] methionine and cysteine (∼10 MBq/mL). When SecG insertion was analyzed, SecA (60 μg/mL) was additionally included. INV was added to the mixture 10 min after the start of the reaction. The translation/insertion reaction was allowed to proceed at 25°C for 90 min. The reaction was then terminated by chilling the mixture in ice. An aliquot (3 μL) was treated with 5% trichloroacetic acid (TCA) to monitor the translation level. The other aliquot (15 μL) was digested with proteinase K (0.5 mg/mL) to generate the membrane-protected fragments (MPF) as an index of membrane insertion, followed by TCA (5%) precipitation. The proteins were then analyzed by SDS-PAGE and autoradiography, as described (57). The radioactive bands were visualized by Phosphorimager (Typhoon, GE). The individual bands were quantified using the ImageQuant software (GE).

### Lipid Extraction and TLC

Purified inner membranes (30 μg) with YibN overexpressed or control (empty vector) were diluted to a final volume of 200 μL in TSG buffer. The diluted membrane was then mixed with 800 μL of methanol: chloroform solution (2:1 vol/vol). After15 min at room temperature, 200 μL of chloroform and 200 μL of dH2O were added sequentially to the samples and mixed briefly. The two phases were separated by centrifugation at 3,000 x rpm for 10 min. The organic phase, which contains lipids, was dried under nitrogen. The dried lipids were resuspended in 30 μL of chloroform and spotted onto a TLC Silica gel (Millipore). The TLC plate was developed in a solution composed of 35:25:3:28 (vol:vol) of chlorofrom:triethylamine:dH2O:ethanol. Plates were air dried and Coomassie stained (58).

### Transmission electron microscopy

A single colony of BL21 bearing pBAD33-YibN or empty pBAD33 plasmid was picked to inoculate in a 10 mL overnight culture in LB media supplemented with 30 µg/mL chloramphenicol. The overnight culture was back-diluted 1/100 into 10 mL fresh LB media supplemented with chloramphenicol and grown until OD600 reached 0.4. Both cultures (BL21 pBAD33 YibN and BL21 pBAD33) were induced with 0.2% arabinose for another 2 hours. Cells were harvested by centrifugation (6,000 x g, 10 minutes). Cell pellets (10 µL) obtained from BL21 pBAD33 YibN and BL21 pBAD33 were treated with a fixative buffer (2.5% glutaraldehyde, 4% formaldehyde, and 0.1 M sodium cacodylate, pH 6.9), then washed with 0.1 M sodium cacodylate buffer. The cells were then fixed with 1% osmium tetroxide and rinsed with water. Staining was performed with 2% uranyl acetate and the cell pellets were rinsed with water and dehydrated by ethanol. The dehydrated cells were transferred and embedded into Epon Resin. After resin polymerization, thin sections were cut and adsorbed onto electron grids. The samples were visualized under Tecnai Spirit TEM.

## Acknowledgement

We are grateful to Dr. Ross Dalbey’s lab for the kind gift of plasmids PC-Lep and Pf3-23Lep, and Dr. rer. nat. Gabriele Deckers-Hebestreit for the plasmid pET20 atpE and the anti-F0c antibody. We acknowledge the UBC Bioimaging Facility (RRID: SCR_021304) for the Tecnai Spirit TEM data.

## Funding and additional information

The work was supported by the Natural Sciences and Engineering Research Council of Canada (DG AWD-008949 to FDvH and DG-20234 to MB), the Canadian Institutes for Health Research (FDN-148408 to CMO), and the Japan Society for the Promotion of Science (KAKENHI 24H01107, 23H04536, 22H02567 to K-i N). CMO is a Canada Research Chair.

